# Identification of regulatory microRNAs for hypoxia induced coagulation mechanism by *In-silico* analysis

**DOI:** 10.1101/2020.06.26.173112

**Authors:** Anju A. Hembrom, Swati Srivastava, Iti Garg, Bhuvnesh Kumar

## Abstract

**Background:** Hypoxia (oxygen deprivation) is known to induce a prothrombotic state by activating the process of coagulation. Hypoxia inducible factor (HIF), is a major transcription factor involved in cellular response to low oxygen tension. MicroRNAs (miRNAs) are non-coding regulatory sequences capable of post-translational modification of multiple genes. Present study aimed to identify common regulatory miRNAs of HIF gene family and coagulation pathway by *In-silico* approach in order to understand the molecular mechanism underlying the process of hypoxia induced coagulation.

**Methods:** HIF family and Coagulation pathway genes along with their receptors and mediators were enlisted after comprehensive literature review. Thorough search of three highly cited database was done to identify the miRNAs targeting these genes. The extracted miRNA list was then prioritized on the basis of number and functional relevance of genes targeted. The target genes of all prioritized miRNAs were listed and subjected to followed by gene ontology (GO) study and pathway analysis.

**Result:** Present *In-silico* analysis identified five candidate miRNAs, viz., hsa-mir-4433a-3p, hsa-mir-4667-5p, hsa-mir-6735-5p, hsa-mir-6777-3p and hsa-mir-6815-3p that co-regulate genes of HIF family and coagulation pathway. GO and pathway analysis revealed that genes regulated by these five microRNAs, predominantly modulate genes facilitating coagulation and hypoxic response.

**Conclusion:** We identified five key microRNAs involved in body’s response to hypoxia and potentially involved in the blood coagulation cascade as well. These five candidate miRNAs may serve as putative epigenetic biomarkers and therapeutic targets to modulate the process of hypoxia induced coagulation.

## Introduction

Hypoxia, as the name suggests is a condition of insufficient oxygen at tissue level, which in turn affects cellular respiration. When the body senses the decline in oxygen availability, it undergoes adaptive changes to overcome this stress. As a coping mechanism of the body, it implies changes in gene expression levels that may either promote survival in oxygen deprivation or enhance oxygen delivery to the cells (1). Hypoxia may arise due to various reasons, like tissue hypoxia at tumor sites, as outcome of hyper-proliferation; any other classical heart or respiratory diseases like asthma, high altitude (HA) hypoxia due to decrease in partial pressure of oxygen, and sometimes even overexertion during exercise or physical labor. Hypoxia Inducible Factor-1 (HIF-1) is a transcription factor, which is a key regulator facilitating the induction of genes for adaptation at hypoxic as well as normoxic conditions (2). HIF-1 is a heterodimer of oxygen dependent HIF-1α and constitutively expressed HIF-1β. The HIF family belongs to a basic Helix-Loop-Helix Per-ARNT-Sim (bHLH-PAS) family of proteins which is required for this dimerization and mainly comprises of 5 members; HIF-1α, HIF-2α (a product of Endothelial PAS1 gene), HIF-3α, ARNT (Aryl Hydrocarbon Nuclear Translocator) also known as HIF-1β and ARNT2 (3). The HIF-3α also has a splice variant IPAS (Inhibitory-PAS) which is dominant negative regulator of HIF-1α and prevents its DNA binding activity (4). The HIF-1 heterodimer translocates into the nucleus and binds to HRE (Hypoxia Response Element) at the promoter region of the gene and regulates its transcription.

Several studies have been done in past to study effect of hypoxia on homeostatsis. During normal homeostatic conditions, the body maintains equilibrium between the pro-coagulants and anti-coagulating factors. However, exposure to hypoxic stress can increase the risk of developing thrombosis by hypoxia-induced platelet aggregation and activation of blood coagulation cascade (5). Hypoxia has been demonstrated to create a pro-thrombotic milieu by an increase in thrombin generation in whole blood even in absence of physical exercise (6). As platelet activation is the primary cause of hemostatsis during vascular injury, substantial evidences demonstrate platelet activation and an increase in platelet adhesiveness in healthy volunteers on exposure to hypoxia (7). The proteomic profiling of hypoxic platelets revealed differentially expressed coagulation proteins like Calpain in accordance to hypoxia exposure at HA along with increased P-selectin levels, the platelet activation biomarker (8); Although, this heightened platelet activity may itself not be the concluding factor for thrombotic precipitation, but accumulated substrate for coagulation may favor this predisposition. These findings augment an explanation as to why venous and not arterial thrombosis is more frequent at HA (9).

Different studies demonstrate increased fibrinogen levels and platelet count with respect to hypoxia, which may further result in clot formation. In addition to that, platelet α-granules are known to store key factors of coagulation pathway, such as, fibrinogen, Factor V, Factor VIII and vWF, fibrinolytic proteins and prominent anticoagulants like tissue factor (TF) and tissue factor pathway inhibitor (TFPI) also, which locally attribute to coagulation (10). Platelet count, which is essentially a contributing factor for platelet aggregation, has been shown to decrease on hypoxia exposure, which may be due to increased platelet consumption for aggregation (7). A longitudinal study has also reported mean platelet volume and plasma fibrinogen levels to increase on ascent and stay at HA (11).

MicroRNAs are epigenetic modulator of gene expression post-transcriptionally regulating the expression of protein-coding genes by non-complimentary base pairing with the mRNA of the target genes affecting their protein levels, without modification in their gene sequences. Since the complimentarity between mRNA and miRNA is imperfect pairing largely depending on the seed sequences (∼8 nucleotides) and not the entire ∼22 nucleotide stretch, a single miRNA can have multiple mRNA targets (12). The fate of the mRNA strand also depends on the degree of complimentarity between the two sequence, as it determines how the later will be silenced, either through decrease in mRNA levels or a translation inhibition (13). MiRNAs have property of altering mRNA expression at post-transcriptional level for number of genes simultaneously. This makes them a strong candidate for target-mediated expressional study, to be used as biomarker or therapeutic target for various complex diseases.

In this study we tried to identify potential miRNAs which co-regulate expression of genes in response hypoxic exposure (HIF family genes) as well as genes of the coagulation cascade. *In-silico i*dentification of regulatory miRNAs of hypoxia and coagulation to understand the common molecular mechanism linking the two, may serve as a future direction for extrapolating and validating it in animal models and cell-culture experiments.

## Methodology

Ethical approval was not necessary for this in-slico analysis as it is a compilation of previously predicted and validated data from different websites.

### Candidate gene selection

This study is focused on the systematic analysis of hypoxia related microRNAs in coagulation. We intended to identify microRNAs that target genes which co-regulate the adaption of the body to hypoxic conditions as well as involved in the process of hemostasis or blood coagulation. Complying with the objective of the study, a list of all the five hypoxia inducible factor (HIF) family genes and genes involved in blood coagulation cascade was prepared (considering only the intrinsic and extrinsic pathway) with a thorough search of the Pubmed and KEGG pathway database (https://www.genome.jp/kegg/pathway.html).

### Data sources and retrieval

For the selected candidate genes, an attempt to retrieve the most miRNA targets was made by searching three reliable and highly cited databases for *in-silico* studies and miRNA target search namely, MiRWalk (http://mirwalk.umm.uni-heidelberg.de/) (14), MiRNet (https://www.mirnet.ca/miRNet/home.xhtml) (15) and miRTargetLink Human (https://ccb-web.cs.uni-saarland.de/mirtargetlink/) (16). A thorough in-depth search of all the three databases was done without setting up any filters to retrieve all the possible targets. A vast data list obtained from different databases was compiled together and was carefully analyzed to filter out any pre-mature miRNAs and repeated entries among different databases.

### Inclusion Criteria

The filtered data obtained was sorted again with inclusion criteria of only those miRNAs that co-regulate both the HIF family genes as well as genes of the coagulation pathway. Implementing this eligibity condition the list was further reduced. Yet there were huge discrepancies among the obtained miRNAs list. To homogenize the data, the miRNA list was sorted on the basis of the number and types of targeted genes, this subsequently allowed to prioritize the miRNAs targeting most genes from both the list.

### Target Identification

As an outcome of the previous step, a list of high priority candidate miRNAs regulating both the HIF family and coagulation genes was obtained. To confirm that these miRNAs do not have a ubiquitous expression and specifically target the genes associated with expression of proteins essential for adaptation to hypoxia, simultaneously contributing to coagulation and hemostasis, a reverse approach was applied. All the target genes of these high priority candidate miRNAs were enlisted with three previously used databases i.e, miRWalk, miRNet and miRTargetLink Human.

### Target Validation

The target gene lists obtained were then subjected to gene ontology analysis and pathway analysis and sorted on the basis of resultant gene ontology results and the selected biological pathways. This was necessary to validate that the target gene list predominantly regulated the hypoxia triggered coagulation cascade and assisted the process of thrombus development by manipulating the major pathways. This sorting was done with the help of online database ShinyGO v0.61 (http://bioinformatics.sdstate.edu/go/) (17) and the Panther database (http://www.pantherdb.org/) (18) respectively.

## Results

This study provides a thorough *in-silico* analysis of the microRNAs co-regulating HIF family genes and the genes of the coagulation cascade and uses a systematic approach to the predict candidate miRNAs which overlap in biological functioning of both the complex regulatory networks. To further check that the genes were not ubiquitously expressed and were within the interests of the aim of the study, we tried to further validate the outcomes of the *in-silico* results with a reverse approach and retrieved the target gene list of the candidate miRNAs and performed their gene ontology and pathway analysis. As an outcome of the study we observed that the major factors affecting coagulation and body’s response to hypoxia were within the target of these predicted candidate miRNAs, infact other complimentary signaling pathways facilitating clot formation like endotheling signaling and plasminogen activation cascade were also predominant targets.

### MiRNAs regulating HIF family genes

HIF family genes were selected namely HIF1α, HIF2α, HIF3α, ARNT and ARNT2. The initial miRNA call for these genes enrolled more than 2300 miRNAs entries. Thorough check for removing pre-mature products, duplicates and sorting out the common regulatory miRNAs was done.

### MiRNAs regulating Coagulation genes

The selection and sorting method of miRNAs was same as that for HIF family genes. Sincere effort was made to include all the coagulation cascade genes including their receptors and mediators with the help of pathway network available online. Some genes had very rare target binding site for miRNAs, i.e., a few entries of regulating miRNAs, however a complete list of all the available predicted and validated miRNA targets from the three databases was extracted and it included all the predicted and validated miRNA targets for forty two coagulation pathway genes.

### Hypoxia related miRNAs regulating Coagulation genes

MiRNAs which had target biding site for both HIF family genes, as well as coagulation cascade genes were sorted out. The miRNAs exclusively regulating either of the two gene families were excluded. After extensive sorting we obtained a list of 1445 miRNA entries. The results obtained were very heterogenous. We further grouped the miRNAs on the basis of the number and types of HIF family genes along with number of coagulation pathway genes being their targets. For selection of candidate miRNA, this list was then prioritized on the basis of most HIF genes regulated. At the end of this comprehensive process we obtained 5 candidate miRNAs; hsa-miR-4433a-3p, hsa-miR-4667-5p, hsa-miR-6735-5p, hsa-miR6777-3p, hsa-miR-5815-3p which targeted all five genes of HIF family and many other genes of coagulation pathway, as depicted in figure 2. Apart from these 5 miRNAs listed, there were other miRNAs which also regulated atleast 4 genes of the HIF family and other coagulation pathway genes, as listed in Table 1.

**Table 1.** Other candidate miRNAs list prioritized on the basis of number and type hypoxia and coagulation pathway genes regulated by them.

| Sl. No. | MicroRNAs | HIF Family Genes | Coagulation Pathway Genes |
| --- | --- | --- | --- |
| 1. | hsa-mir-5698<br>hsa-mir-4633-3p<br>hsa-mir-504-5p<br>hsa-mir-6782-5p<br>hsa-mir-6893-5p | ARNT, ARNT2,<br>HIF1A, HIF3A | F2, F3, F8, F9, F11, F2R, F2RL1, F2RL2, F2RL3,<br>FGB, PLAT, PLAUR, PROCR, TFPI, PLG, PLGLB1,<br>PLGLB2, THBD, CPB2, KNG1, KLKB1, SERPINA1,<br>SERPINE1 |
| 2. | hsa-mir-6510-5p | ARNT, HIF1A,<br>HIF2A, HIF3A | F13B, F2RL2, F5, F8, F9, FGA, FGB, KNG1, PLGLB1,<br>PLGLB2, PROCR, PROS1, SERPINA1, SERPINE1 |
| 3. | hsa-mir-3157-5p | ARNT2, HIF1A,<br>HIF2A, HIF3A | F2RL1, PLGLB1, PLGLB2, PROS1 |
| 4. | hsa-mir-4701-3p | ARNT, ARNT2,<br>HIF1A, HIF2A | F13A1, F5, FGB, PLAT, PLGLB1, PLGLB2, PROCR,<br>SERPINA1, TFPI |

**Figure 1.**
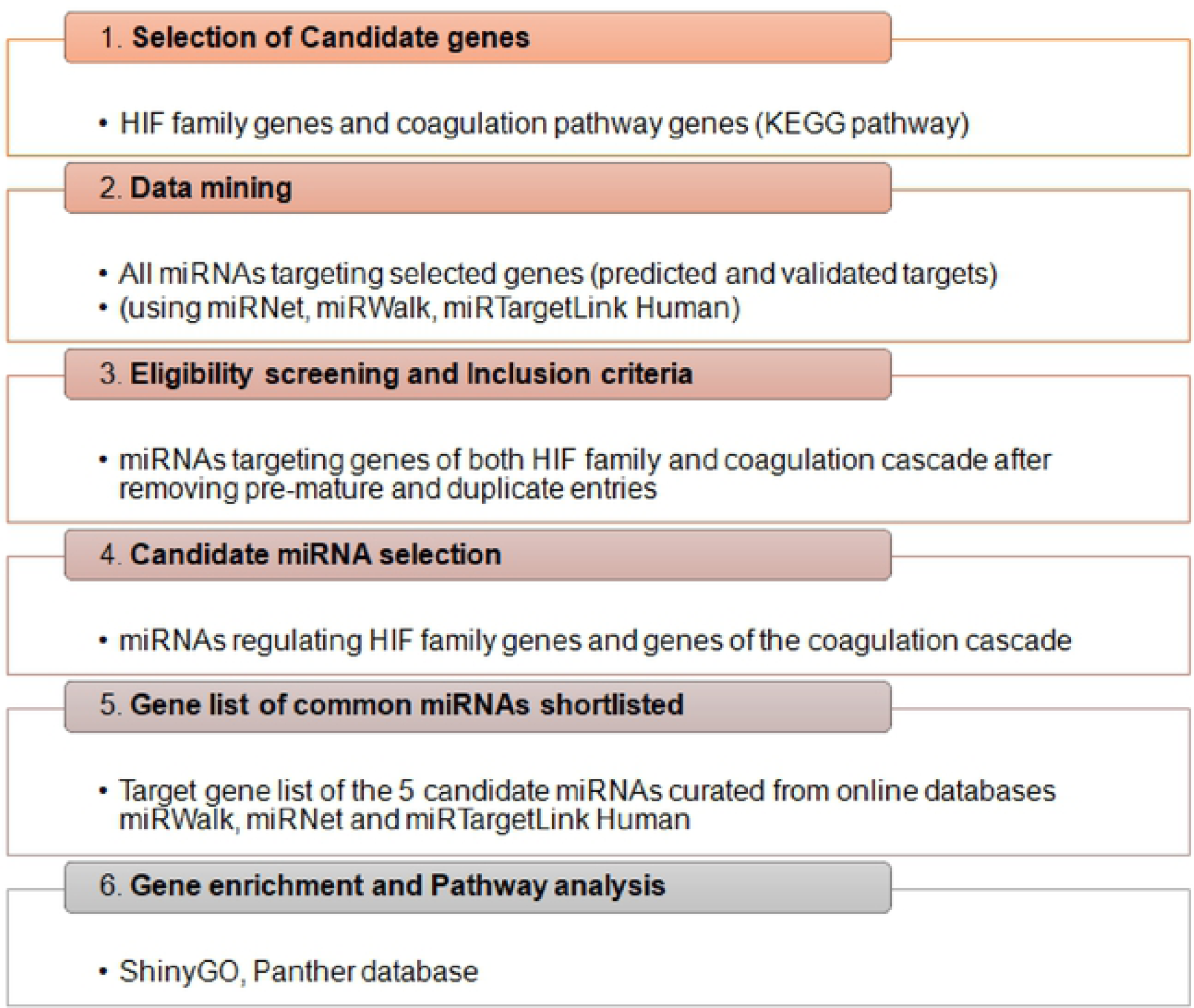
Diagramatic representation for the study work-flow

**Figure 2.**
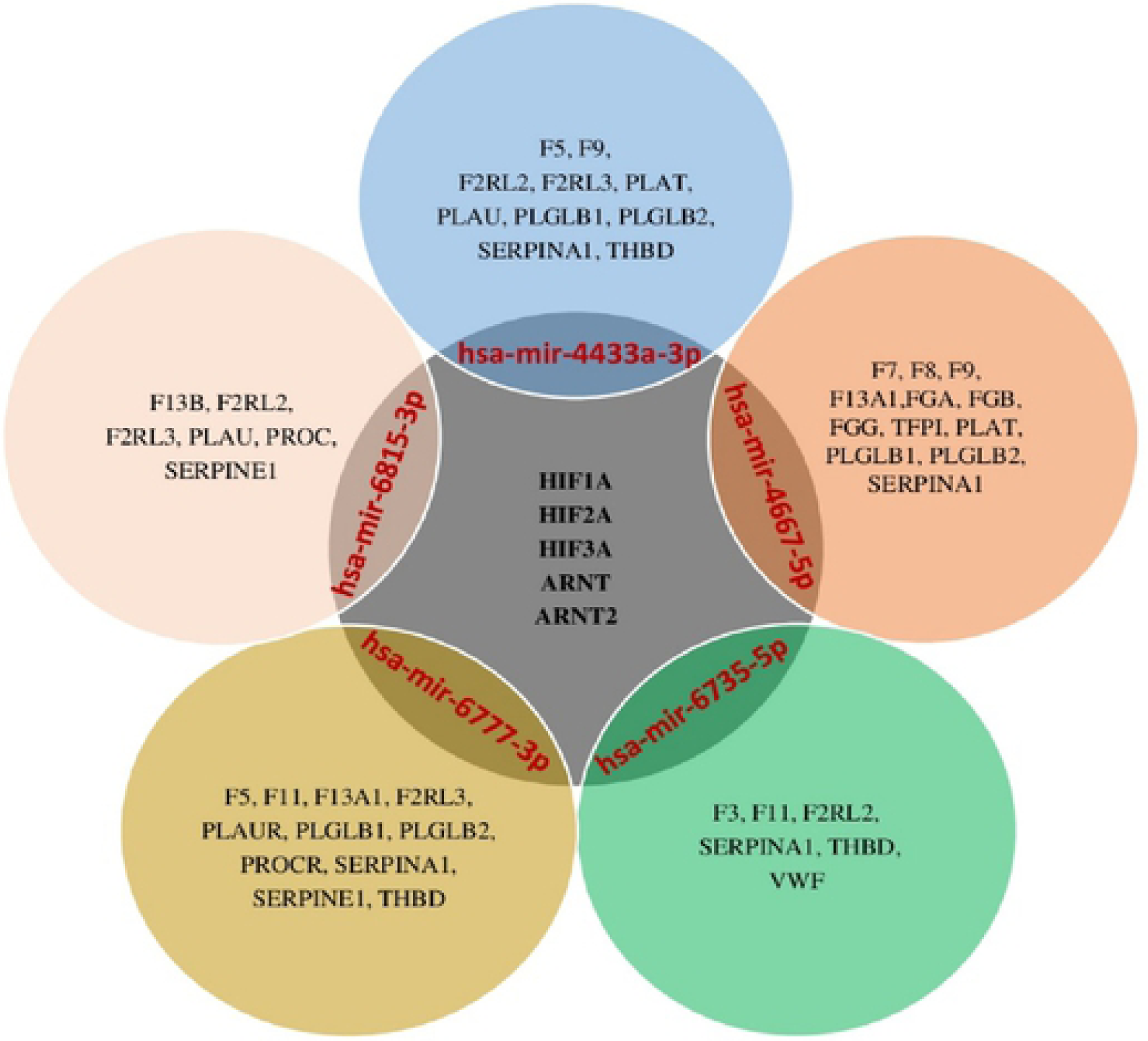
Diagrammatic representation of the 5 candidate microRNAs (hsa-mir-4433a-3p, hsa-mir-4667-5p, hsa-mir-6735-5p, hsa-mir-6777-3p, and hsa-mir-6815-3p) that regulate genes of hypoxia as well as coagulation pathway.

### Gene Ontology and Biological Pathway targets

The target genes of prioritized miRNAs when subjected to gene ontology analysis and pathway analysis which yielded a list of significant biological, cellular and molecular processes as well as pathways affected. An insight into pathway analysis further elaborated a wide diversity of target signaling cascades, which included major pathways affecting blood coagulation and thrombus formation with a special emphasis on hypoxia response via HIF activation, plasminogen activation cascade, and PDGF signaling. Other important signaling pathways like oxidative stress response, EGF and FGF receptor signaling and endothelin signaling pathways were also regulated. Molecular functions indicated that bonding, transcription regulator and catalytic activities were targeted. Similarly, cellular components like cell, membrane, organelle, and protein containing complexes were also regulated (figure 3A).

**Figure 3.**
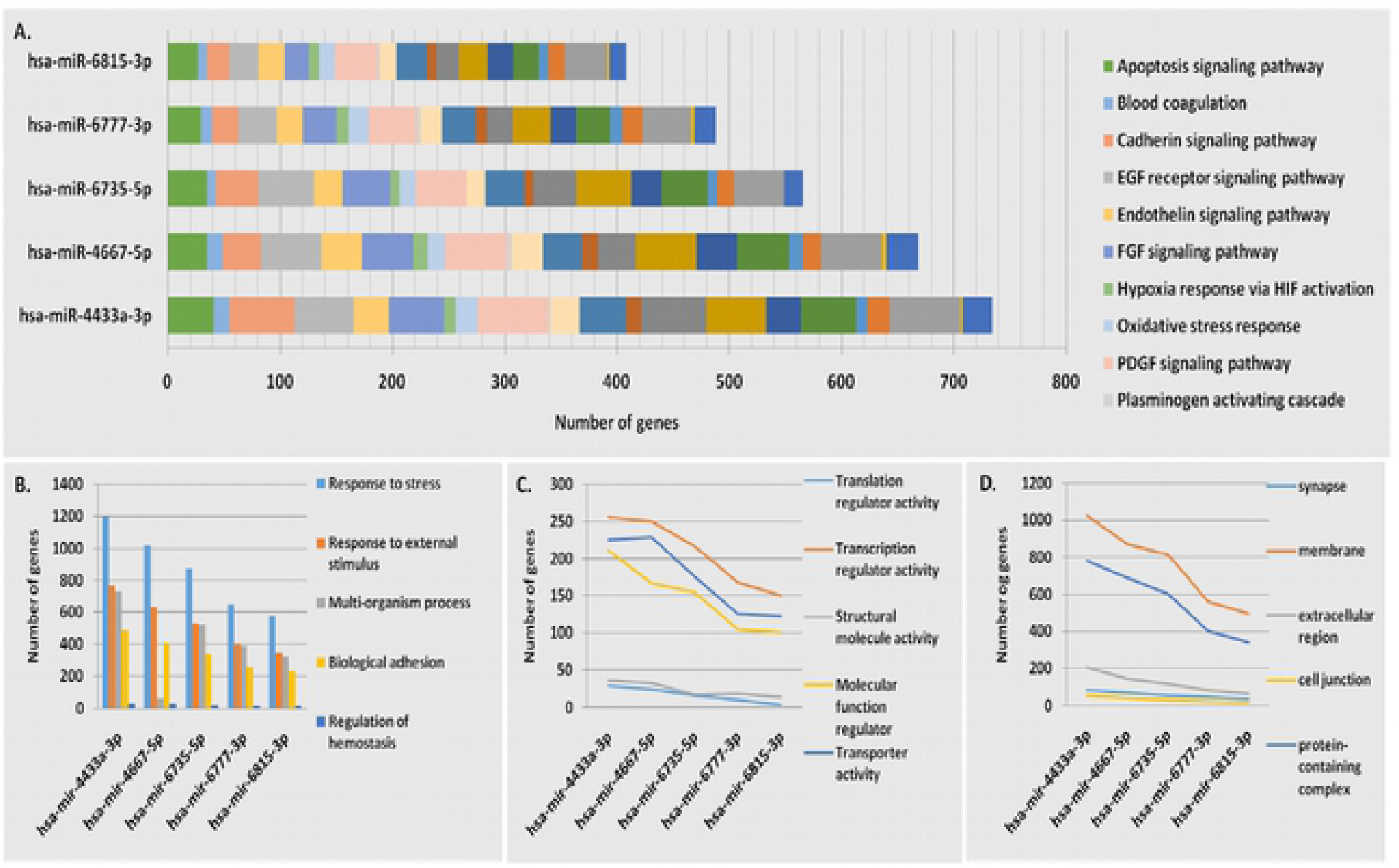
Gene Ontology and biological pathway targets results using ShinyGO database showing (A. Biological pathways) and Panther database showing (B. Biological process, C. Molecular functions, D. Cellular components).

Amongst significant biological processes were response to stress, response to external stimuli, biological adhesion, regulation of hemostasis etc. (figure 3B). Predominant molecular functions included transcription and translation regulator, structural molecule activity, transporter activity etc (figure 3B). Similarly, cellular components like cell junction, membrane, extracellular region and protein containing complexes were also regulated (figure 3C).

## Discussion

During conditions of physiological stress, such as hypoxia, the body is more susceptible to succumb to pro-thrombotic state like those at high altitudes. Hypoxia acts as a trigger for activating the genes of coagulation cascade, regardless of the physical activity (19). Various studies have shown that hypoxia predisposes the homeostasis shift towards a pro-coagulative milieu. When this equilibrium is disturbed; blood clotting disorders like stroke, deep vein thrombosis, pulmonary embolism are likely to occur. Hypobaric hypoxic environment prevailing at HA results in increase in the viscosity of blood due to increase in pro-coagulatory factors (20), such as platelet aggregation, rise in fibrinogen levels and polycythemia, which further leads to thrombotic state (21). It is an undeniable fact that HIF family genes regulate a surfeit of pathological symptoms in humans (22), however, they are responsible for adaptation to hypoxia. Since microRNAs are the key modulators and master regulators of gene expression, they are considerably important in regulating the expression of HIF regulated genes. In an attempt to establish a common regulatory link between coagulation and hypoxia, the findings of this systematic *in-silico* analysis meticulously consolidates miRNAs which overlap in function by regulating genes of both the hypoxia and as well as coagulation cascade. With the help of a thorough search of highly cited databases like miRWalk, mirTargetLink Human and miRNet, the present study yielded a list of five miRNAs, hsa-mir-4433a-3p, hsa-mir-4667-5p, hsa-mir-6735-5p, hsa-mir-6777-3p and hsa-mir-6815-3p. These five miRNAs targeted all five genes of HIF gene family along with several genes of coagulation cascade. To further validate these findings, a reverse approach was used to find all the target genes of shortlisted miRNAs followed by gene ontology search and pathway analysis. The outcome corroborated with our results, as the genes of shortlisted miRNAs were not ubiquitously expressed and regulated major biological processes like regulation of hemostasis, biological response to stress, and biological adhesion. They were shown to regulate biological pathways like hypoxia response via HIF activation along with blood coagulation, PDGF signaling and endothelin signaling; thus predominantly facilitating activation of hypoxia induced coagulation cascade. In addition to this, a second priority list of miRNAs was also prepared (Table 1) which target four genes of HIF gene family and various other genes of coagulation pathway. This study is a first of its kind to contemplate a thorough and curated list of priority miRNAs which may be used as a biomarker or even as a therapeutic target in regulating coagulopathy under hypoxia environment. These miRNAs could serve as a reference to extrapolate the findings of this study in cell lines and animal models and implicate the roles of these predicated candidate miRNAs as biomarker or therapeutic targets.

## Conclusion

*In silico* analysis using various bio-informatics tools narrow down to five miRNAs as most promising candidates for determining hypoxia induced coagulation mechanism. These miRNAs could be used as potential biomarkers or therapeutic targets that could be modulated for regulating hypoxia activated coaulopathy.

## Funding Received

The author(s) received no specific funding for this work.

## Acknowledgements

Authors are extremely grateful to technical staff and research scholars of our group for logistic support

